# Boundary cells in the representation of episodes in the human hippocampus

**DOI:** 10.1101/2021.05.28.446233

**Authors:** Hye Bin Yoo, Gray Umbach, Bradley C. Lega

## Abstract

The representation of episodes is a fundamental requirement for forming episodic memories, but the specific electrophysiological mechanisms supporting episode construction in the human hippocampus remain unknown. Experiments in rodent models indicate that a population of neurons sensitive to edges of an environment, termed *border* or *boundary* neurons in spatial navigation, fulfills a role analogous to episode demarcation. We hypothesized that such boundary neurons could be identified in the human mesial temporal lobe, with firing rates sensitive specifically to the beginning and end of mnemonically-relevant episodes in the free recall task. Using a generalized linear model to control for factors such as encoding success and item onset times along with other variables, we found 44 *Boundary* neurons out of a total 736 single neurons recorded across 27 subjects. We distinguish boundary neurons from a separate population of ramping neurons, which are time-sensitive neurons whose activity provides complementary but distinct information during episodic representation. We also describe evidence that the firing of boundary neurons within the preferred windows (at the beginning and end of episodes) is organized by hippocampal theta oscillations, using spike-field coherence metrics.

## Main

A key feature of episodic memory is the ability to construct distinct episodes out of continuous experience (Howard et al., 2012). Episode construction requires demarcation of when an episode begins and ends, facilitating item associations within these temporal boundaries (Clewett et al., 2019). Behavioral evidence indicates that the boundaries create a discontinuity in the temporal associations of encoded items (Ezzyat & Davachi, 2011), promote the clustering of events by relative contexts (DuBrow & Davachi, 2013), and affect the temporal structure of retrieved memories (Heusser et al., 2018). The electrophysiological mechanisms of boundary construction constitute a critical question in human neuroscience. A direct analogy to episode demarcation may be the representation of boundaries in space, a function supported by border neurons or boundary neurons (Barry et al., 2006; Savelli et al., 2008; Solstad et al., 2008), which exhibit sensitivity of firing rate to geometric boundaries. Based on the hypothesized similarity between spatial and temporal contextual representations (see Eichenbaum, 2017), these data predict that temporal analogues of border neurons may demarcate *episodic* boundaries. Preliminary evidence for boundary-like MTL activity has come from human subjects viewing movie scenes (Zheng et al., 2021). Another class of MTL neurons that may participate in boundary construction is ramping neurons (Tsao et al., 2018). Ramping cells exhibit logarithmic decreases (or increases) in firing rate relative to the beginning or end of groups of events across different time scales (Umbach et al., 2020). However, whether ramping neurons co-exist with boundary neurons and how their properties differ remain unknown. We sought to find a population of temporal *boundary* neurons that are distinct from *ramping* neurons as subjects performed an episodic memory task. We identified both classes of neurons using activity recorded from microelectrodes implanted in human MTL.

The microelectrode recordings and initial processing used in this study were previously described in Umbach et al., 2020. Twenty-seven human epilepsy patients with implanted intracranial micro-electrodes for seizure recording at Thomas Jefferson University Hospital (TJ) or University of Texas Southwestern (UT) participated in the study. The IRBs from both institutions approved this study. A total of 40 recording sessions were collected using Behnke-Fried style microelectrodes (Ad-Tech, Oak Creek, WI). Identification and isolation of individual units utilized Combinato (Niediek et al., 2016), with results directly comparable to other studies in humans (Faraut et al., 2018; Umbach et al., 2020). Specifics are reported in Detailed Methods.

Participants performed a free recall episodic memory task, consisting of between four and 25 lists comprised of 12 or 15 memory items (common nouns) followed by a math distractor task and then a 30- or 45-second retrieval period during which participants freely recalled as many items as possible. During the encoding period for each list, subjects were given a sequence of words on a laptop screen that each lasted for 1.6 seconds. Each word was temporally separated by a jittered gap ranging from 0.8 to 1.2 seconds. In the distractor period, subjects typed in answers to simple arithmetic problems (A + B + C = ?), where A, B, and C were random nonzero one-digit integers.

We defined Boundary and Ramping cells using a generalized linear model (GLM) based identification routine motivated by previous studies (Reddy et al., 2020; Tsao et al., 2018; Umbach et al., 2020). First, a continuous time series representing probabilistic firing rate was constructed per neuron by applying a Gaussian kernel function on the spike train whose values are one at the time a spike is detected. The firing rate curve was incorporated into a GLM as the dependent variable. We selected independent variables as: 1) boundary for encoding and retrieval epochs in free recall, 2) ramping (positive or negative direction corresponding to up or down ramping) during task-relevant epochs, 3) item onset of encoded words regardless of recall status, 4) onset of successfully encoded words, 5) vocalization at retrieval, and 6) resting or inactive task condition between completion of a retrieval epoch and the subsequent encoding epoch. The first two were the predictors of interest in modeling, whereas the rest were control predictors for excluding neurons responding to these other factors (most importantly, recall success). We used *stepwiseglm* with log-link function (MATLAB 2019b, The MathWorks Inc, Natick, MA) to model the firing rate curve assuming an exponential relationship between predictors and the activity. The model selected relevant independent variables based on the goodness-of-fit estimated by *R*^2^ so that if Δ*R*^2^ was larger than 0.01 the model included the predictor, but removed those with Δ*R*^2^ lower than 0.005. Further specifics are shown in Detailed Methods.

We required the following three conditions for the definition of a Boundary or Ramping cell: 1) the neuron’s firing rate should be modeled significantly by the final model that includes either the boundary or ramping predictor (but not both), 2) the magnitude of *t* score of boundary or ramping should be the highest among all included predictors, and 3) log-likelihood for the unrestricted model that includes all predictors should be significantly greater than a restricted model excluding only boundary or ramping predictor depending on the neuron type of interest (MATLAB’s *lratiotest, df* = 1, *p* < 0.05). Additionally for Boundary cells, only those with a positive model coefficient (U-shape) firing rate changes were included. Boundary or Ramping cell populations were mutually exclusive based on these requirements. As a result, we separately identified 44 Boundary (6%) and 75 Ramping (10%) neurons out of a total 736 single units. The proportion of Boundary cells was significantly smaller than Ramping (*Z* test, *Z* = −3.777, *p* < 0.001). Out of 40 sessions, 11 contributed at least one Boundary cell, and 18 contributed at least one Ramping cell. Figure 1 shows two sample neurons, and their normalized firing rate curves averaged across all encoding and retrieval periods. The resulting curves are consistent with expectations based on the modeling criteria, i.e. a Boundary cell exhibits an asymmetric U-shaped curve, while a Ramping cell exhibits an increase across the epoch. Figure 2a shows that the average activity curve from all Boundary and Ramping cells demonstrate the expected pattern of activity during encoding and retrieval. We emphasize that Boundary cell activity does not reflect memory success effects (namely, primacy and recency in the free recall task) as neurons responding to encoding success separate from boundary conditions are explicitly excluded based on the parameters of the GLM.

**Figure 1:**
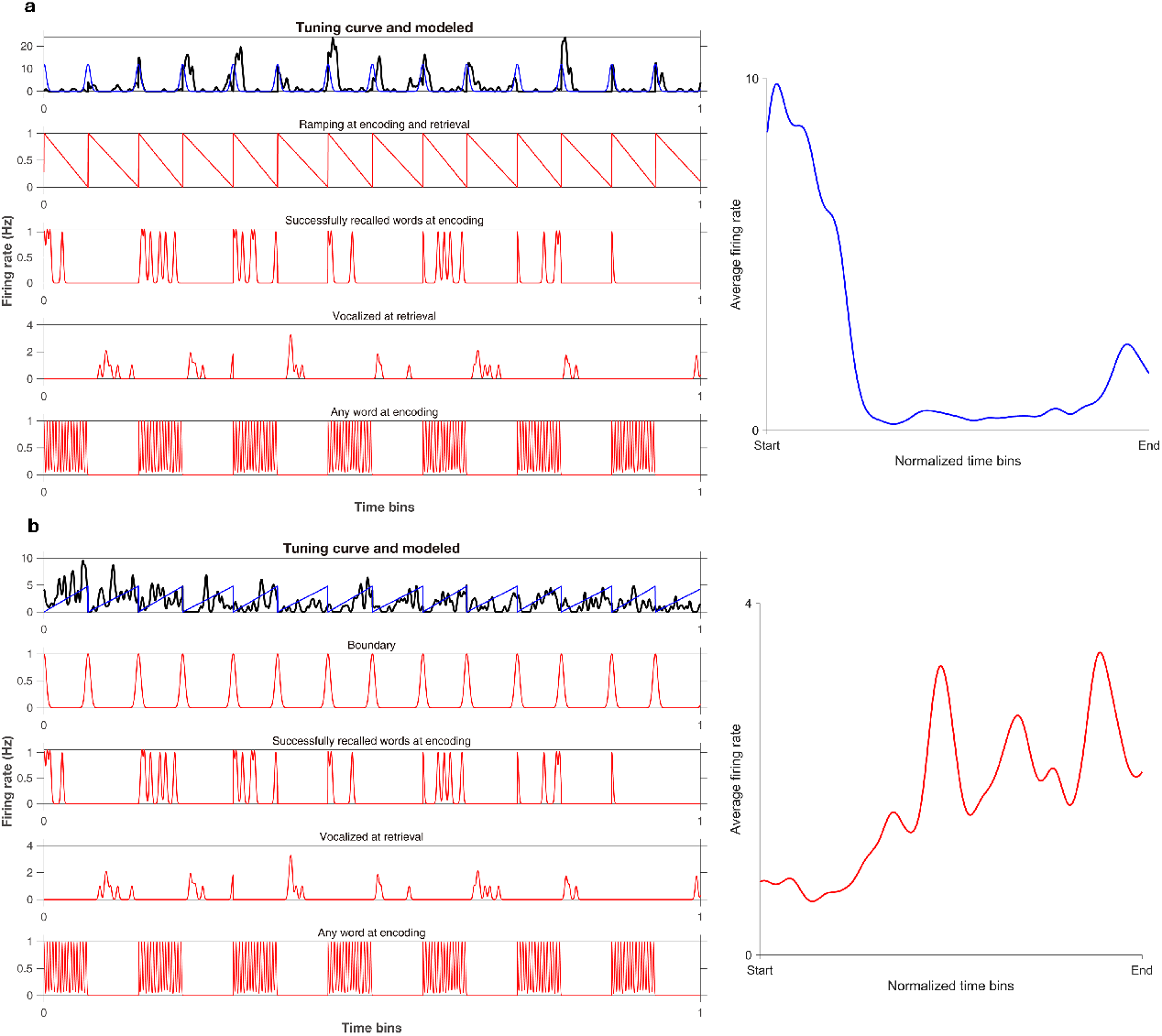
Characteristics of sample Boundary and Ramping cells. **a**, Activity (black) of a sample Boundary cell modeled by predictors of interest (blue) on the top row, excluding the effect of control predictors (red). Activity curve averaged across all encoding and retrieval conditions of the sample Boundary cell is demonstrated on the right. **b**, Activity (black) of a sample Ramping cell modeled by predictors of interest (blue) on the top row, excluding the effect of control predictors (red). Activity curve averaged across all encoding and retrieval conditions of the sample Ramping cell on the right.

**Figure 2:**
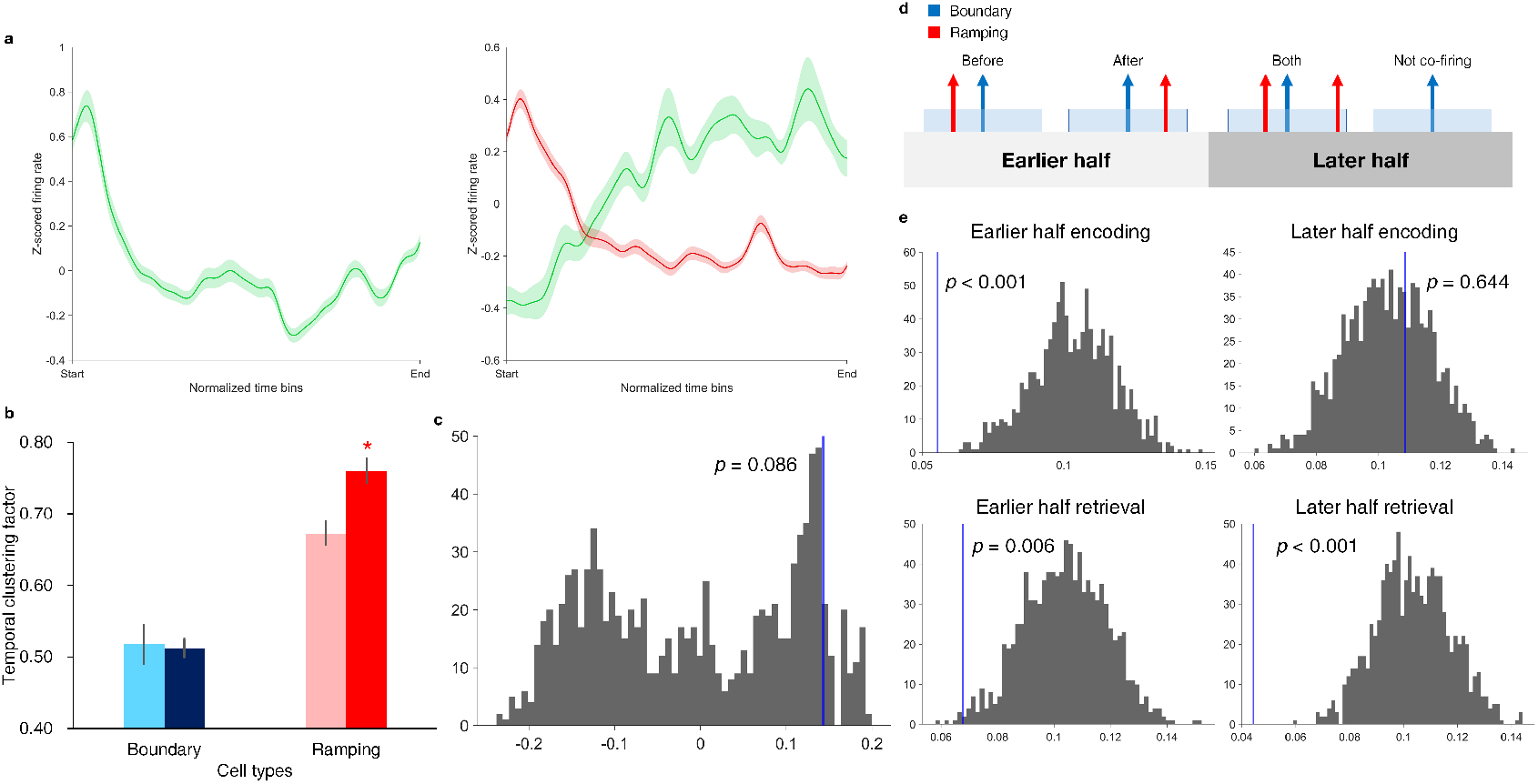
Boundary and Ramping cells’ characteristics, behavioral associations and the co-firing of two groups. **a**, Activity curve averaged across all encoding and retrieval conditions of Boundary (left) and Ramping (right) cells. Shades denote SEM. Green denotes average of neurons that have positive model coefficients, and red negative coefficients. **b**, Comparison of temporal clustering behavior associated with Boundary (*blue*) and Ramping (*red*) cells demonstrating median-split higher-(darker) and lower-(lighter) magnitude model coefficients. Bar height represents the mean and error bars the SEM. Star indicates a significant difference (*p* < 0.05) via rank-sum test. **c**, Permutation testing comparing the correlations between the model coefficient and temporal clustering factor for Ramping than Boundary cells. **d**, Representation of three cases where any Ramping cells co-fire within ± 25 ms of a Boundary cell spike, which are labeled *Before, After* and *Both*, and a null case where no co-firing occurs. Four time windows of interest were considered, namely the earlier and later halves of encoding and retrieval periods. **e**, Permutation tests comparing the real versus random medians of co-firing fractions. Co-firing was more sparse than chance except for the later half of encoding.

We performed a permutation test to confirm the robustness of Boundary and Ramping detection via the GLMs. For each neuron, spike times were circularly randomized maintaining their gap lengths to create 1,000 random firing rate curves. The same independent variables modeled the randomized curves. We compared the ratios of positive calls (true positive + false positive) over the total (736) for Boundary and Ramping models and compared them against the actual likelihood ratios (44/736, 75/736 respectively). Permutation test showed that the actual likelihood ratio is significantly higher than the randomized positive likelihood ratio (*p* < 0.001 for both groups), confirming that the actual fraction of Boundary and Ramping found using model definitions is significant over chance.

We related the model coefficients of Boundary and Ramping cells with behavior using 1) performance of free recall, 2) successful recall ratio of the first and last (boundary) items on lists within the free recall task, and 3) temporal clustering factor values (TCF), which quantify the tendency of recalling contiguously presented items during retrieval (Howard & Kahana, 2002; Manning et al., 2012; Polyn et al., 2009; Umbach et al., 2020). We tested for correlations between the magnitude of *t* scores from Boundary and Ramping models and these three behavioral scores observed from sessions corresponding to each neuron, using a median split applied to the magnitude of model-derived *t* scores, with a rank-sum test to compare the behavioral scores from the higher versus lower *t* score groups. Boundary model *t* scores did not significantly relate to any behavioral score (*p >* 0.182). However, Ramping model *t* scores predicted the magnitude of temporal clustering (*p* < 0.001) and boundary recall (*p* = 0.037). Figure 2b represents the result of comparing TCF between lower and higher model coefficient groups of Boundary and Ramping cells using rank-sum test. Non-parametric Spearman’s correlation confirmed that higher Ramping *t* scores correlate with higher TCF (*r* = 0.491, *p* < 0.001), and higher boundary recall (*r* = 0.326, *p* = 0.004). Correlation with TCF remained significant when incorporating overall performance or boundary recall success into the predictive model (partial correlation, *r* = 0.453, *p* < 0.001, *r* = 0.430, *p* < 0.001, respectively). This finding for Ramping cells is consistent with our previously published findings related to time sensitive cells in the MTL (Umbach et al., 2020). When we compared these two populations (Boundary vs Ramping cells) directly using the median difference in temporal clustering score, we observed a trend towards significance (Figure 2c, *p* = 0.086). These findings provide preliminary evidence that Ramping but not Boundary cell firing provides the information necessary for making temporal associations in the free recall task. We discuss some possible interpretations of these findings below.

We examined whether Boundary and Ramping cells exhibit co-firing. This analysis was motivated first by the question of whether Boundary cells represent an integration of Ramping cell activity, a model that would entail the expectation of significant co-firing during task. The time scale we selected (25 ms) over which to test for co-firing was motivated by the findings of Harris et al., 2003, indicating that this specific scale is highly relevant for the construction of neuronal assemblies in associative memory formation. Our analysis of co-firing was necessarily limited to (six) sessions in which both Boundary and Ramping cells were identified. To maximize sensitivity, co-firing instances were separately counted for the earlier and later halves of the entire encoding and retrieval periods. A graphical illustration of co-firing analysis is shown in Figure 2d. As such, we tested for co-firing during four temporal epochs: earlier and later *encoding*, and *retrieval*. Per-mutation testing using randomized spike times was performed to compare the proportion of real co-firing over chance. We found that the Boundary cells co-fire with Ramping cells significantly *less* than chance throughout encoding and retrieval (*p* ≤ 0.006), except during the second half of encoding lists (*p* = 0.644, Figure 2e). At minimum, these findings indicate that MTL Boundary cell firing cannot be explained directly as the integration of Ramping cell activity, although the relative scarcity of Boundary cells (with few simultaneous Boundary/Ramping sessions available for analysis) limits our conclusions in this regard. However, our observations have some support in findings from rodents during spatial navigation, addressed below in the Discussion section.

Phase locking relative to hippocampal theta oscillations may be a mechanism for integrating Boundary neurons’ activity with other features of episode representation. We therefore hypothesized that phase locking would be greater for spikes occurring at boundaries for Boundary cells, as compared to non-boundary windows. Thus, we tested for theta (<10 Hz) phase locking in Boundary cells via spike-field coherence (SFC) (Fries et al., 2001; Fries et al., 1997; Rutishauser et al., 2010), to test if they fire more in-phase at boundaries than non-boundary windows. We calculated SFC for all spikes in and out of boundary windows per neuron (3 seconds, see Detailed Methods). Boundary cells that have at least ten spikes within boundary windows were counted for the calculation, and the number of sampled spikes in and out of boundary windows were equalized (via random downsampling) to avoid biasing the results based on spike frequency. SFC in and out of boundary windows was compared using a permutation test for paired groups per frequency bin, with Holm-Bonferroni correction (Holm, 1979). Figure 3 demonstrates that for slower theta (<5 Hz), Boundary cells exhibit a significant field coherence in boundary versus non-boundary windows (* *correctedp* < 0.01). We also tested for phase precession in Boundary cells, motivated by the properties of place cells (Mehta et al., 2002; O’Keefe & Burgess, 2005) and our previous report on phase precession of time cells (Umbach et al., 2020). Precession was measured following the approach of Kempter et al., 2012. We did not find evidence of phase precession for Boundary cells, as only *n* ≤ 4 out of 44 Boundary cells exhibited a significant circular-linear relationship between spike phase and time within these firing windows. This is somewhat unsurprising, as phase locking and phase precession are complementary mechanisms for organization of spiking activity relative to theta phase.

**Figure 3:**
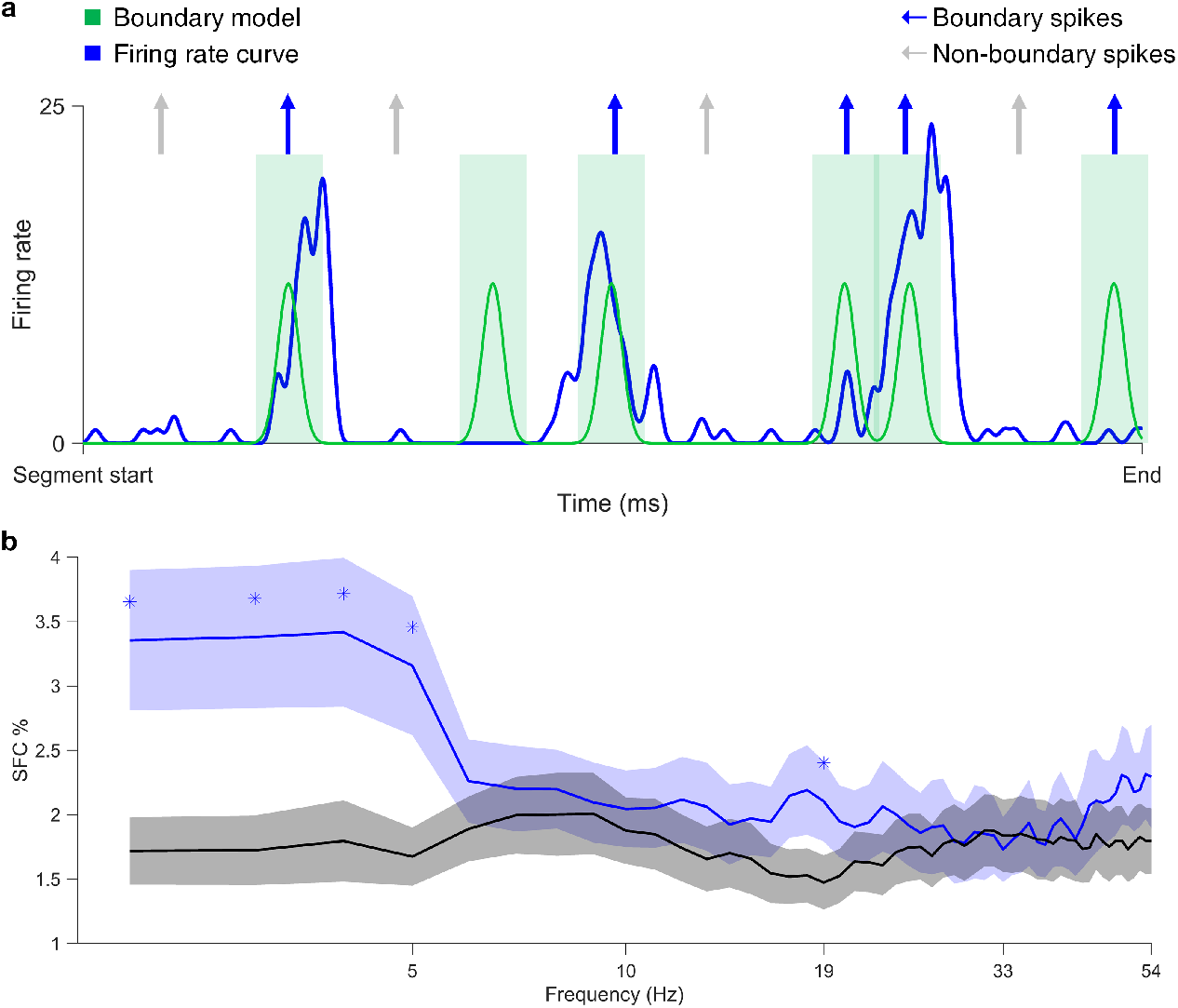
Significant spike-field coherence occurs in boundary compared to non-boundary windows. **a**, A schematic description of selecting boundary and non-boundary spikes out of a spike train. A sample segment of firing rate curve from a Boundary cell (solid blue) is shown superposed on its boundary model predictor (solid green) that mark boundary windows around the beginning and the end of encoding and retrieval periods (green shade). Arrows indicate some boundary (blue) and non-boundary (gray) spikes that are potentially included in the SFC analysis. **b**, Spike-field coherence for boundary windows in Boundary cells is compared against non-boundary windows in the same group of neurons using permutation tests. Stars indicate p-values that are corrected with Holm-Bonferroni method and lower than 0.01. Frequency log-spaced in 1–54 Hz is demonstrated for visualization.

This study demonstrates the existence of unique Boundary cells that represent the demarcation of events in an episodic memory task, using a GLM-based method to eliminate the effects of item onset or recall success. The identification of Boundary cells in the MTL helps explicate the electro-physiological mechanisms supporting episodic memory, and their properties have the potential to inform models of mnemonic processing. Boundary cells may provide an “anchor” signal to promote context reinstatement in models such as the temporal context model (Alexander, Robinson, et al., 2020; Hinman et al., 2019; Julian et al., 2018). Boundary signals further establish important parallels between spatial navigation in rodent models and episodic associations in humans (Alexander, Robinson, et al., 2020; Barry et al., 2006; Horner et al., 2016; van Wijngaarden et al., 2020; Zheng et al., 2021). The significant theta phase locking among Boundary cells specifically during boundary representation may provide a direct mechanism for integration of episodic information with other populations in the MTL and cortex, potentially incorporating inter-regional phase amplitude coupling (D. X. Wang et al., 2021).

Our data propose important questions regarding the information provided by Boundary cells that will require further investigation. First, we found that Boundary cells are relatively less frequent in the MTL as compared to Ramping cells (and *time* cells, see Umbach et al., 2020), which may indicate that Boundary cell activity in the MTL reflects sparse coding of a more detailed boundary/episode representation occurring elsewhere in the cortex. Our findings have support in existing rodent data, in which border-type cells were statistically sparser than ramping cells and are more prominent outside the hippocampus (Alexander, Carstensen, et al., 2020; Bjerknes et al., 2014; C. Wang et al., 2018; Zheng et al., 2021), such as in rhinal (Gofman et al., 2019) and retrosplenial cortex (Alexander, Carstensen, et al., 2020). Boundary cell activity in regions such as the retrosplenial cortex may reflect the integration of cognitive goals and sensory information necessary to determine boundary moments and construct episodes (Alexander, Carstensen, et al., 2020; Barry et al., 2006; Ezzyat & Davachi, 2011) – such extensive information may not be processed directly in the MTL. By contrast, the detailed temporal information from time sensitive cells (ramping and time cells, which occur more frequently in the MTL) suggests that time within an episode may be represented more directly within the MTL (Umbach et al., 2020). This distinction may reflect the proposed difference between more allocentric representations in the MTL, and egocentric representations occurring in regions outside the MTL (Alexander, Carstensen, et al., 2020; Bicanski & Burgess, 2020; Gofman et al., 2019).

A distinction in the type of information represented by Boundary versus Ramping cells is also suggested by a lack of co-firing on short time scales in our data. This observation has some support from rodent findings, in which Bjerknes et al., 2014 showed that border neurons mature earlier than grid cells (a proposed spatial memory analogue of Ramping cells) and contribute differently to spatial memory. We acknowledge, however, that one must be cautious not to over-interpret null results, because only six experimental sessions included both time cells and ramping cells, and the identified Boundary and Ramping cells were both in the MTL. Certainly, one would predict that two cell groups contribute cooperatively to episode representations. A rodent model shows that boundary representation in retrosplenial cortex is at least partially driven by the upstream allocentric information from the MTL (van Wijngaarden et al., 2020), and grid cells are known to change their mapping depending on environment represented by boundaries (Barry et al., 2007). The mechanism of Boundary and Ramping cell integration, perhaps using theta time scales, will ultimately require further investigation, potentially combining microelectrode data across regions.

The lack of a significant behavioral association between Boundary cell firing and temporal clustering, or recall fraction, may be explained by the fact that episode construction is a fundamental requirement of task participation. In other words, the absence of a boundary signal may only occur in patients incapable of understanding the task structure who do not participate in memory testing, and as such a link with memory behavior is not apparent in our task paradigm. A behavioral association for boundary signals may be more readily discernible using experimental paradigms that make behavioral demands on transitions among episodes, as suggested by Zheng et al., 2021. We note that these preliminary findings echo our own results in which boundary activity was a negative predictor of temporal clustering behavior. Such alternate paradigms may further explicate whether Boundary cells can fill a hypothesized role as an “anchor” signal in episodic representations (Alexander, Robinson, et al., 2020), as boundary information can theoretically provide an internal cue for guiding the “tracing” characteristic of episodic representations (Alexander, Robinson, et al., 2020; Bicanski & Burgess, 2020; Bicanski & Burgess, 2018). However, our own data do not test this idea, since the free recall paradigm cannot identify Boundary cell activity that is unique for different contexts. We also note that the specific firing characteristics of the Boundary cells we observed, with asymmetrical firing between beginning versus end of temporal epochs, may further inform how models of episodic memory account for boundary information in delineating episodes.

## Detailed Methods

### Single cell separations

Before spike detection and sorting, We filtered the LFP for broadband noise using a volume conduction subtraction algorithm (Kota et al., n.d.). Using Combinato, we applied a band-pass filter to the raw LFP at 300-1000 Hz for threshold crossing (spike identification), then extracted spike features filters at 300-3000 Hz (Niediek et al., 2016). We inspected: i) the shape of the mean spike waveform; ii) the fraction of inter-spike intervals shorter than 3 ms; iii) the shape of the distribution of inter-spike intervals; iv) the stationarity of unit spiking; and v) similarity to other mean spike waveforms (Faraut et al., 2018). We separated 736 single neurons, and their spike trains were aligned with the corresponding source microwire’s LFP time series data and downsampled to 1000 Hz (1 kHz).

### Design of generalized linear model

We defined the dependent variable of neuronal model using a probabilistic firing rate curve. It was constructed from a neuron’s spike train referring to Baranauskas et al., 2012, as shown below:

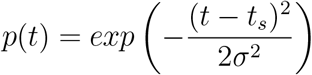

In this equation, *p*(*t*) represents the probability of observing spikes or firing rate (Hz), *t* current time, *t*_*s*_ the spike time, *σ* corresponds to the kernel width in our study, fit to 1 second or 1000 ms for the unit of Hz.

Each independent variable was defined as follows. Boundary in active task conditions referred to the beginning and the end of encoding and retrieval conditions. The distractor condition is automatically accounted for by the boundaries of these two conditions. The beginning and the end of the encoding period is defined as the first word onset and 1.5 seconds after the last word onset, respectively. Retrieval boundaries are defined at the retrieval onset (beginning) and the end, which are at 30 or 45 seconds after the onset. To mitigate the edge effect at the end in case encoding and retrieval are close, we subtracted one second margin of error at the retrieval end. At each boundary we assigned a value of one, and applied the Gaussian kernel function with *σ* = 2000 (ms). Ramping was modeled as a linear increase of probability from 0 to 1 across encoding, distractor and retrieval periods. All encoded words (*σ* = 500), the successfully encoded words (*σ* = 1000), and vocalizations during retrieval (*σ* = 1000) were all assigned a value of one at the onset of each event, and the Gaussian kernel function with designated *σ* values was applied. The resting or inactive task condition was modeled as uniform one between retrieval and the encoding of the next list. Because the result of modeling is dependent on the definition of predictors, we confirmed that the Boundary and Ramping cells are significantly categorized over chance with a permutation test using a randomized spike times and the same predictors multiple times against the real neurons.

### Detection of preferred theta frequency

For precession and phase locking analyses, we focused on theta frequency that is divided into slower (2–5 Hz) and faster (5–9 Hz) ranges based on previous studies showed that two sub–bands have functional dissociation in hippocampus (Goyal et al., 2020; B. C. Lega et al., 2012; B. Lega et al., 2016). We defined a preferred theta for both ranges by calculating the oscillatory frequency. We modeled the actual power spectrum of the encoding period (starting at the first encoded item, ending at 1.5 seconds post the last encoded item) averaged across all lists to a function *A* ∗ *f* ^*α*^ referring to Milstein et al., 2009. The power spectra were calculated using Welch’s method (Welch, 1967) with a 1 kHz sampling rate at each of the log-spaced frequencies for the encoding periods of all lists. We first calculated the power difference of the original against the modeled, then selected the frequency within slower or faster band that showed the maximum elevation of actual power compared to the modeled. Spike phase values were extracted using wavelet transform for each neuron’s representative frequency.

### Spike-field coherence

Spike-field coherence (SFC) is a measure of periodic timing relationship between spikes and the background oscillation independent of power spectrum of the LFP as a function of frequency, as described in Fries et al., 2001. This is represented as the percentage (0–100%) of the oscillation power triggered by spikes above the power averaged across all windows around spikes per frequency bins. A higher SFC indicates that spikes follow a particular phase at that frequency band (Fries et al., 2001; Rutishauser et al., 2010). To calculate SFC, we acquired spike-triggered average (STA) and spike-triggered power (STP). We took a 500 ms long time window of the LFP before and after spike time (total 1.001 seconds), downsampled the signal by the factor of four reducing the sampling frequency to 250 Hz, and obtained the STA by averaging the time series across all windows. We quantified STP by first taking the power spectrum using the multitaper method using Chronux toolbox (Jarvis & Mitra, 2001; Mitra, 2007), at 250 Hz (frequency resolution = 4 Hz) with time-bandwidth product of four and seven tapers, following Rutishauser et al., 2010. The power spectrum from each window was averaged to obtain STP. SFC was the percentage of the power spectrum of STA over STP for each frequency bin covered by the multitaper method (<125 Hz). In this study, we calculated SFC counting spikes in and out of boundary windows, which are defined as 1.5 seconds before and after a boundary epoch marking the beginning and the end of encoding, and retrieval periods (total four per list). To account for the shorter boundary windows (3.001 secs) in comparison to the outside windows, we first counted all spikes in boundary windows and randomly downsampled spikes from non-boundary windows to equalize spike sample sizes.

## Data Availability

Please contact the corresponding author, Bradley Lega for the access to the data and codes implemented for this study.

## Acknowledgements

We sincerely appreciate the subjects volunteered for this study. We are grateful to Dr. Michael Sperling and Dr. Ash Sharan of Thomas Jefferson University Hospital for their work in data collection.

## Notes

**Conflict of Interest** The authors report no conflict of interest.

### Competing Interest Statement

The authors have declared no competing interest.

## References

Alexander, A. S., Carstensen, L. C., Hinman, J. R., Raudies, F., Chapman, G. W., & Hasselmo, M. E. (2020). Egocentric boundary vector tuning of the retrosplenial cortex. Science advances, 6 (8), eaaz2322.

Alexander, A. S., Robinson, J. C., Dannenberg, H., Kinsky, N. R., Levy, S. J., Mau, W., Chapman, G. W., Sullivan, D. W., & Hasselmo, M. E. (2020). Neurophysiological coding of space and time in the hippocampus, entorhinal cortex, and retrosplenial cortex. Brain and Neuroscience Advances, 4, 2398212820972871.

Baranauskas, G., Maggiolini, E., Vato, A., Angotzi, G., Bonfanti, A., Zambra, G., Spinelli, A., & Fadiga, L. (2012). Origins of 1/f2 scaling in the power spectrum of intracortical local field potential. J Neurophysiol, 107 (3), 984–94. https://doi.org/10.1152/jn.00470.2011

Barry, C., Lever, C., Hayman, R., Hartley, T., Burton, S., O’Keefe, J., Jeffery, K., & Burgess, N. (2006). The boundary vector cell model of place cell firing and spatial memory. Rev Neurosci, 17 (1-2), 71–97. https://doi.org/10.1515/revneuro.2006.17.1-2.71

Barry, C., Hayman, R., Burgess, N., & Jeffery, K. J. (2007). Experience-dependent rescaling of entorhinal grids. Nature neuroscience, 10 (6), 682–684.

Bicanski, A., & Burgess, N. (2020). Neuronal vector coding in spatial cognition. Nat Rev Neurosci, 21 (9), 453–470. https://doi.org/10.1038/s41583-020-0336-9

Bicanski, A., & Burgess, N. (2018). A neural-level model of spatial memory and imagery. Elife, 7, e33752.

Bjerknes, T. L., Moser, E. I., & Moser, M.-B. (2014). Representation of geometric borders in the developing rat. Neuron, 82 (1), 71–78.

Clewett, D., DuBrow, S., & Davachi, L. (2019). Transcending time in the brain: How event memories are constructed from experience. Hippocampus, 29 (3), 162–183. https://doi.org/10.1002/hipo.23074

DuBrow, S., & Davachi, L. (2013). The influence of context boundaries on memory for the sequential order of events. Journal of Experimental Psychology: General, 142 (4), 1277–1286. https://doi.org/10.1037/a0034024

Eichenbaum, H. (2017). On the integration of space, time, and memory. Neuron, 95 (5), 1007–1018. https://doi.org/10.1016/j.neuron.2017.06.036

Ezzyat, Y., & Davachi, L. (2011). What constitutes an episode in episodic memory? Psychological science, 22 (2), 243–252.

Faraut, M. C. M., Carlson, A. A., Sullivan, S., Tudusciuc, O., Ross, I., Reed, C. M., Chung, J. M., Mamelak, A. N., & Rutishauser, U. (2018). Dataset of human medial temporal lobe single neuron activity during declarative memory encoding and recognition. Sci Data, 5, 180010. https://doi.org/10.1038/sdata.2018.10

Fries, P., Reynolds, J. H., Rorie, A. E., & Desimone, R. (2001). Modulation of oscillatory neuronal synchronization by selective visual attention. Science, 291 (5508), 1560–1563.

Fries, P., Roelfsema, P. R., Engel, A. K., König, P., & Singer, W. (1997). Synchronization of oscillatory responses in visual cortex correlates with perception in interocular rivalry. Proceedings of the National Academy of Sciences, 94 (23), 12699–12704.

Gofman, X., Tocker, G., Weiss, S., Boccara, C. N., Lu, L., Moser, M.-B., Moser, E. I., Morris, G., & Derdikman, D. (2019). Dissociation between postrhinal cortex and downstream parahippocampal regions in the representation of egocentric boundaries. Current Biology, 29 (16), 2751–2757. e4.

Goyal, A., Miller, J., Qasim, S. E., Watrous, A. J., Zhang, H., Stein, J. M., Inman, C. S., Gross, R. E., Willie, J. T., Lega, B., Lin, J. J., Sharan, A., Wu, C., Sperling, M. R., Sheth, S. A., McKhann, G. M., Smith, E. H., Schevon, C., & Jacobs, J. (2020). Functionally distinct high and low theta oscillations in the human hippocampus. Nat Commun, 11 (1), 2469. https://doi.org/10.1038/s41467-020-15670-6

Harris, K. D., Csicsvari, J., Hirase, H., Dragoi, G., & Buzsaki, G. (2003). Organization of cell assemblies in the hippocampus. Nature, 424 (6948), 552–6. https://doi.org/10.1038/nature01834

Heusser, A. C., Ezzyat, Y., Shiff, I., & Davachi, L. (2018). Perceptual boundaries cause mnemonic trade-offs between local boundary processing and across-trial associative binding. Journal of Experimental Psychology: Learning, Memory, and Cognition, 44 (7), 1075.

Hinman, J. R., Chapman, G. W., & Hasselmo, M. E. (2019). Neuronal representation of environmental boundaries in egocentric coordinates. Nature communications, 10 (1), 1–8.

Holm, S. (1979). A simple sequentially rejective multiple test procedure. Scandinavian journal of statistics, 65–70.

Horner, A. J., Bisby, J. A., Wang, A., Bogus, K., & Burgess, N. (2016). The role of spatial boundaries in shaping long-term event representations. Cognition, 154, 151–164.

Howard, M. W., Viskontas, I. V., Shankar, K. H., & Fried, I. (2012). Ensembles of human mtl neurons “jump back in time” in response to a repeated stimulus. Hippocampus, 22 (9), 1833– 1847. https://doi.org/10.1002/hipo.22018

Howard, M. W., & Kahana, M. J. (2002). A distributed representation of temporal context. Journal of Mathematical Psychology, 46 (3), 269–299.

Jarvis, M. R., & Mitra, P. P. (2001). Sampling properties of the spectrum and coherency of sequences of action potentials. Neural Comput, 13 (4), 717–49. https://doi.org/10.1162/089976601300014312

Julian, J. B., Keinath, A. T., Frazzetta, G., & Epstein, R. A. (2018). Human entorhinal cortex represents visual space using a boundary-anchored grid. Nature neuroscience, 21 (2), 191– 194.

Kempter, R., Leibold, C., Buzsaki, G., Diba, K., & Schmidt, R. (2012). Quantifying circular-linear associations: Hippocampal phase precession. J Neurosci Methods, 207 (1), 113–24. https://doi.org/10.1016/j.jneumeth.2012.03.007

Kota, S., du Plessis, A., Massaro, A. N., Chang, T., Al-Shargabi, T., & Govindan, R. B. (n.d.). A frequency based spatial filter to mitigate volume conduction in electroencephalogram signals, In 2016 38th annual international conference of the ieee engineering in medicine and biology society (embc), IEEE.

Lega, B. C., Jacobs, J., & Kahana, M. (2012). Human hippocampal theta oscillations and the formation of episodic memories. Hippocampus, 22 (4), 748–61. https://doi.org/10.1002/hipo.20937

Lega, B., Burke, J., Jacobs, J., & Kahana, M. J. (2016). Slow-theta-to-gamma phase-amplitude coupling in human hippocampus supports the formation of new episodic memories. Cereb Cortex, 26 (1), 268–278. https://doi.org/10.1093/cercor/bhu232

Manning, J. R., Sperling, M. R., Sharan, A., Rosenberg, E. A., & Kahana, M. J. (2012). Spontaneously reactivated patterns in frontal and temporal lobe predict semantic clustering during memory search. J Neurosci, 32 (26), 8871–8. https://doi.org/10.1523/JNEUROSCI.5321-11.2012

Mehta, M. R., Lee, A. K., & Wilson, M. A. (2002). Role of experience and oscillations in transforming a rate code into a temporal code. Nature, 417 (6890), 741–6. https://doi.org/10.1038/nature00807

Milstein, J., Mormann, F., Fried, I., & Koch, C. (2009). Neuronal shot noise and brownian 1/f(2) behavior in the local field potential. Plos One, 4 (2), e4338. https://doi.org/ARTNe433810.1371/journal.pone.0004338

Mitra, P. (2007). Observed brain dynamics. Oxford University Press.

Niediek, J., Bostrom, J., Elger, C. E., & Mormann, F. (2016). Reliable analysis of single-unit recordings from the human brain under noisy conditions: Tracking neurons over hours. PLoS One, 11 (12), e0166598. https://doi.org/10.1371/journal.pone.0166598

O’Keefe, J., & Burgess, N. (2005). Dual phase and rate coding in hippocampal place cells: Theoretical significance and relationship to entorhinal grid cells. Hippocampus, 15 (7), 853–66. https://doi.org/10.1002/hipo.20115

Polyn, S. M., Norman, K. A., & Kahana, M. J. (2009). A context maintenance and retrieval model of organizational processes in free recall. Psychol Rev, 116 (1), 129–56. https://doi.org/10.1037/a0014420

Reddy, L., Zoefel, B., Possel, J., Peters, J., Dijksterhuis, D., Poncet, M., van Straaten, E. C., Baayen, J. C., Idema, S., & Self, M. W. (2020). Human hippocampal neurons track moments in a sequence of events. bioRxiv.

Rutishauser, U., Ross, I. B., Mamelak, A. N., & Schuman, E. M. (2010). Human memory strength is predicted by theta-frequency phase-locking of single neurons. Nature, 464 (7290), 903–7. https://doi.org/10.1038/nature08860

Savelli, F., Yoganarasimha, D., & Knierim, J. J. (2008). Influence of boundary removal on the spatial representations of the medial entorhinal cortex. Hippocampus, 18 (12), 1270–1282.

Solstad, T., Boccara, C. N., Kropff, E., Moser, M. B., & Moser, E. I. (2008). Representation of geometric borders in the entorhinal cortex. Science, 322 (5909), 1865–8. https://doi.org/10.1126/science.1166466

Tsao, A., Sugar, J., Lu, L., Wang, C., Knierim, J. J., Moser, M. B., & Moser, E. I. (2018). Integrating time from experience in the lateral entorhinal cortex. Nature, 561 (7721), 57–62. https://doi.org/10.1038/s41586-018-0459-6

Umbach, G., Kantak, P., Jacobs, J., Kahana, M., Pfeiffer, B. E., Sperling, M., & Lega, B. (2020). Time cells in the human hippocampus and entorhinal cortex support episodic memory. Proc Natl Acad Sci U S A, 117 (45), 28463–28474. https://doi.org/10.1073/pnas.2013250117

van Wijngaarden, J. B., Babl, S. S., & Ito, H. T. (2020). Entorhinal-retrosplenial circuits for allocentric-egocentric transformation of boundary coding. Elife, 9, e59816.

Wang, C., Chen, X., Lee, H., Deshmukh, S. S., Yoganarasimha, D., Savelli, F., & Knierim, J. J. (2018). Egocentric coding of external items in the lateral entorhinal cortex. Science, 362 (6417), 945–949.

Wang, D. X., Schmitt, K., Seger, S., Davila, C. E., & Lega, B. C. (2021). Cross-regional phase amplitude coupling supports the encoding of episodic memories. Hippocampus, 31 (5), 481– 492. https://doi.org/10.1002/hipo.23309

Welch, P. D. (1967). The use of fast fourier transform for the estimation of power spectra: A method based on time averaging over short, modified periodograms. IEEE Transactions on audio and electroacoustics, 15 (2), 70–73.

Zheng, J., Gómez Palacio Schjetnan, A., Yebra, M., Mosher, C., Kalia, S., Valiante, T. A., Mamelak, A. N., Kreiman, G., & Rutishauser, U. (2021). Cognitive boundary signals in the human medial temporal lobe shape episodic memory representation. bioRxiv, 2021.01.16.426538. https://doi.org/10.1101/2021.01.16.426538

